# Incidence and attributes of chimeric COI and 18S sequences derived from nematode-infected arthropods

**DOI:** 10.1101/2025.05.16.654383

**Authors:** Jacopo D’Ercole, Robin Floyd, Sean WJ Prosser, Saeideh Jafarpour, Paul DN Hebert

## Abstract

High-throughput sequencing is speeding the discovery of new species and making it possible to quantify the species richness of bulk samples. However, chimeric sequences can lead to both type I errors (true species mistakenly rejected) and type II errors (chimeras incorrectly viewed as valid species). This study employed nanopore sequencing to examine the incidence and nature of chimeric sequences recovered from 531 arthropod specimens infected with nematodes. Specifically, it examined chimera formation in two gene regions: the 658 bp segment of mitochondrial COI employed as the barcode region for the animal kingdom, and a 900 bp segment of nuclear 18S often employed to discriminate lineages of nematodes. This work revealed chimeras for both gene regions despite the deep sequence divergences between members of these two phyla. However, the incidence of chimeric molecules was higher for 18S than COI, with chimera formation correlating with local variation in DNA stability and conserved regions between parent sequences. Aside from demonstrating that chimeras can arise from distantly related organisms, this study provides insights into the mechanisms underlying their formation.

## Introduction

DNA barcoding has been widely adopted for specimen identification (Hebert et al., 2003) and species discovery (Hebert et al., 2004; Burns et al., 2008). These same features also make it a powerful tool for describing (D’Ercole et al., 2022; Dapporto et al., 2022; D’Ercole et al., 2024; Dapporto et al., 2024) and monitoring (Hardulak et al., 2020; Van Der Heyde et al., 2020; Rodríguez-Ezpeleta et al., 2021) genetic diversity. High-throughput sequencing (HTS) has advanced the application of DNA-based identification systems by making it possible to ascertain the species composition of bulk and environmental samples (i.e., metabarcoding and eDNA) (Cruaud et al., 2017; Liu et al., 2017; Prosser et al., 2025), and to reveal species interactions (i.e., symbiome) (Minardi et al., 2022; Bass et al., 2021). However, the detection of rare species in both contexts is complicated because the misinterpretation of sequence data can create both type I errors (where valid species are mistakenly rejected) and type II errors (where sequencing artifacts are viewed as true species).

Chimeric amplicons are prime candidates for misinterpretation because they combine partial sequences from two or more species. Despite this, their incidence, structural properties, and impact on DNA barcoding/metabarcoding have seen little investigation although the mechanisms underlying their formation is understood. Most chimeras arise when the elongation of an amplicon terminates prematurely, creating a truncated product which can serve as a primer in subsequent PCR cycles. When an incomplete product from one species binds to template from a second taxon and extension proceeds, it produces an interspecific chimeric sequence (Wang and Wang, 1996; Thompson et al., 2002; Haas et al., 2011). Their presence creates two primary issues. First, chimeric sequences inhibit the amplification of target template molecules. Functioning as DNA templates themselves, they compete for reagents throughout amplification, reducing PCR efficiency (Kalle et al., 2014). Second, chimeric sequences are distinct from both their parent taxa, and if not recognized, can be misinterpreted as representing new species (Acinas et al., 2004; Ashelford et al., 2006). Therefore, it is important to characterize chimeric molecules and identify the factors that influence their formation.

This study examines sequence variation in two gene segments commonly used to assess species diversity, the barcode region of mitochondrial cytochrome *c* oxidase subunit I (COI) and the nuclear ribosomal 18S rRNA. It examines the incidence and structure of chimeric molecules created when DNA extracts derived from nematode-infected arthropods are PCR amplified.

Although no primers are universal, those for COI (Hebert et al., 2018) and 18S (Floyd et al., 2002) target highly conserved regions so they are likely to amplify templates from both the arthropods and nematodes, meeting the basic requirement to generate chimeric sequences.

## Methods

### Study design

The Centre for Biodiversity Genomics has used the Sequel and Sequel II platforms (Pacific Biosciences) to sequence amplicons of the 658 bp barcode region of COI from more than 10 million DNA extracts, each derived from a single arthropod. Inspection of the sequence arrays from 0.1 million of these specimens revealed 531 cases where both arthropod and nematode sequences were recovered, indicating that the arthropod carried a nematode infection. These DNA extracts provided a perfect opportunity to investigate the incidence of chimeric molecules in the amplicon pool from each specimen. A pairwise comparison of the arthropod and nematode sequences from these 531 specimens revealed an average divergence of 35% for COI and 24% for 18S. Given this high divergence, chimeras between the arthropod and nematode sequences would be easily recognizable.

### Sampling and genetic characterization

The 531 arthropods included 527 insects and 4 arachnids. The 658 bp barcode region of COI and approximately 900 bp of 18S were amplified with Platinum Taq (Thermo Scientific) using standard primers: C_LepFolF/C_LepFolR (Hebert et al., 2018) for COI and SSU_F_07/SSU_R_26 (Floyd et al., 2002) for 18S. Primers were asymmetrically indexed with unique molecular identifiers (UMIs) to allow the pooling of PCR products from all 531 specimens in a single sequencing run. Primer and UMI sequences are provided in Supplementary table 1.

Each PCR included: 6.25 μl of 10% trehalose (Fluka Analytical), 1.62 μl of molecular grade water (Hyclone), 1.25 μl of 10X Platinum Taq buffer (Thermo Scientific), 0.625 μl of 50 mM MgCl_2_ (Thermo Scientific), 0.0625 μl of 10 mM dNTPs (KAPA Biosystems), 0.625 μl of 2 μM forward primer (IDT), 0.0125 μl of 100 μM reverse primer (IDT), 0.06 μl of 5 U/μl Platinum Taq (Thermo Scientific), and 2 μl of DNA for a total reaction volume of 12.5 μl. PCR for COI employed 45 cycles with the following regime: 94°C for 2 min; 5 cycles of 94°C for 40 sec, 45°C for 40 sec, 72°C for 1 min; 40 cycles of 94°C for 40 sec, 51°C for 40 sec, 72°C for 1 min; 72°C for 2 min. PCR for 18S employed 35 cycles with the following regime: 94°C for 2 min; 35 cycles of 94°C for 40 sec, 56°C for 40 sec, 72°C for 1 min; 72°C for 5 min.

Library preparation for both genes followed ONT recommendations. Briefly, each amplicon pool underwent end preparation using the NEBNext Ultra™ II End Repair/dA-Tailing Module (New England Biolabs, NEB), followed by purification with AMPure-XP beads. Next, ONT sequencing adapters (SQK-LSK114) were ligated onto the amplicons using NEB Quick T4 DNA Ligase (NEB), followed by a final purification step with AMPure-XP beads. Sequencing was performed on a Mk1b ONT sequencer using either a MinION (FLO-MIN114, version 10.4.1) or a Flongle (FLO-FLG114, version 10.4.1) flow cell. The COI library was analyzed on a single MinION flow cell, while the 18S library was analyzed on two Flongles which were combined before further analysis. The higher read depth provided by the MinION flow cell was essential for COI, as a preliminary analysis of the Sequel data only detected 27 COI chimeric sequences across the 531 samples (data not shown), highlighting the need for increased sequencing depth.

### Bioinformatic pipeline for high-throughput sequencing

Sequence analysis was performed in an Ubuntu environment (22.04.3 LTS) employing the informatics pipeline outlined by Hebert et al. (2024). This pipeline included base calling, demultiplexing, filtering reads based on length of the target amplicon, and clustering reads into OTUs. The clustering process entails two stages: an initial round delineated operational taxonomic units (OTUs), while the second reduced the impact of PCR and sequencing errors. To further mitigate the latter errors, OTUs were only retained if they were supported by five or more reads. The VSEARCH SINTAX algorithm (Rognes et al., 2016) was used for taxonomic assignment with a minimum probability threshold of 60%. The MinION flow cell generated 6,930,263 reads, of which 2,645,405 were retained after filtering. By comparison, the two Flongles produced 1,071,419 reads, with 187,183 retained after filtering. The full COI dataset derived from the MinION flow cell was employed for comprehensive examination of the chimeras derived from COI amplicons. However, to enable a direct comparison of the incidence of chimeras between COI and 18S, the MinION dataset was haphazardly downsized to match the read count achieved by the two Flongles. UCHIME (Edgar et al., 2011; Edgar et al., 2016) was employed for initial chimera detection using the de novo approach, which does not rely on a chimera-free reference dataset. This was followed by a manual inspection involving three steps. First, for each sample containing chimeric sequences, FASTA files were aligned using MAFFT (Katoh et al., 2002). Second, when more than two potential parent sequences were identified, only the two with the highest read counts were considered as parents. Third, the analysis focused exclusively on chimeras with a single crossover event, as these can be unambiguously identified and they represent the majority of all chimeras (Zhang et al., 2010).

### Statistical analysis

Histograms displaying sequence recovery for the two gene regions, along with those illustrating the distribution of sequences per sample, were generated using the “ggplot2” (Wickham, 2016) (ver. 3.5.3) package in R Studio.

Hartigan’s Dip Test (Hartigan and Hartigan 1985) was performed and using the “diptest” (ver. 0.77.1) package on R Studio to assess the unimodality of the distribution of midpoint values of crossover ranges. This was followed by the application of a Kernel density estimate, using the package “KernSmooth” (ver. 2.23.26) to create a smooth representation of the density. This integrated approach of statistical testing and visualization offered a robust method for evaluating the unimodal characteristics of the dataset.

A custom R script (Supplementary Script 1) was used to visualize the crossover range of the chimeric molecules characterized by a single crossover region. Chimeras were grouped according to the origin of their parent sequences. The parent sequences were classified into three categories (Unknown, Same, Mix,) based on whether they originated from unknown phyla, same phylum, or different phyla. The graphical representation was generated using the package “ggplot2” (ver. 3.5.2) in R Studio.

Analysis of variance (ANOVA) was conducted using R Studio to assess whether order-level taxonomy influences the mid-point and width of crossover regions. Additionally, ANOVA was applied to determine if the origin of the parent sequences affects the mid-point and width of the crossover regions.

Distinct majority consensus sequences were identified from three separate datasets: one comprised of the parent sequences that generated the COI chimeric molecules in the full COI dataset, one for the parent sequences identified in the reduced COI dataset, and the other consisting of the parent sequences in the 18S dataset. A custom R script was then employed to analyze variation in GC content (Supplementary script 2) and DNA stability (ΔG°) (Supplementary script 3) along these consensus sequences using a sliding window approach. This analysis is important because high GC content and DNA stability hinders the ability of DNA polymerase to extend strands, creating potential hotspots for the formation of chimeras. This analysis employed a window size of 30 base pairs and a step size of 1 base pair. ΔG° values in this analysis considered both base pairing (which relies on the hydrogen bonds between opposite nucleotides) and base stacking (which depends on the interactions between neighboring nucleotides on the same strand) (Santa Lucia, 1998). Trends in ΔG° were modeled and visualized using a Generalized Additive Model (GAM) framework implemented with the R package “mgcv” (Wood, 2011) (ver. 1.9.1). GAM makes it possible to capture non-linear relationships between predictors and the response variable without assuming a specific relationship between the parameters.

A sliding window analysis with a window size of 5 base pairs and a step size of 1 base pair was used to determine if crossover regions in chimeric molecules coincided with regions of high sequence similarity between the parent molecules (Supplementary script 4). This approach was chosen because partially extended DNA strands require a few nucleotides of complementary sequence at their 3’ ends for efficient binding to a different template (Robertson and Walsh-Weller, 1998). To examine this, the percentage identity was determined for three datasets: one containing the parent sequences that produced the COI chimeric molecules in the full COI dataset, one with parent sequences that produced chimeras in the reduced COI dataset, and the other including parent sequences that generated the 18S chimeric molecules. A GAM function was employed to model the trends in percentage identity for both gene regions.

### Data Access

The 531 host arthropod records, along with associated metadata (images, geographic coordinates), are available in the public dataset “DS-CHIMERA” (dx.doi.org/10.5883/DS-CHIMERA).

## Results

### Comparison of COI and 18S

This comparison examined the number of chimeras detected in the 18S and downsized COI dataset. For each genetic marker, 187,183 reads were analysed. The COI and 18S sequences for all 531 samples, along with their corresponding taxonomic assignments, are provided in the .xlsx file of Supplementary table 2.

### Sequence recovery

The comparison of sequence distribution across the 531 samples revealed a difference between COI and 18S. COI was recovered from 522 of the 531 specimens with 199 (37.5%) producing two OTUs, 300 (56.5%) producing 3 or more OTUs, and 23 (4.3%) delivering just one OTU. By comparison, 18S was recovered from 518 specimens with 273 (51.4%) producing two OTUs, 165 (31.1%) producing three or more OTUs, and 80 (15.1%) producing a single OTU (Supplementary figure 1).

COI analysis generated OTUs from 506 of the 531 arthropods (95.3%), and nematode sequences from 460 of them (86.6%). From this total, 444 specimens (83.6%) generated both arthropod and nematode sequences, 62 (11.7%) only included arthropod sequences, while 16 (3%) only delivered nematode sequences (Figure 1A). 18S analysis generated sequences from 322 arthropods (60.6%) and from 468 nematodes (88.1%). Overall, 282 specimens (53.1%) generated both arthropod and nematode sequences, while 40 (7.5 %) only had arthropod sequences, and 186 records (35%) only delivered nematode sequences (Figure 1B).

**Figure 1.**
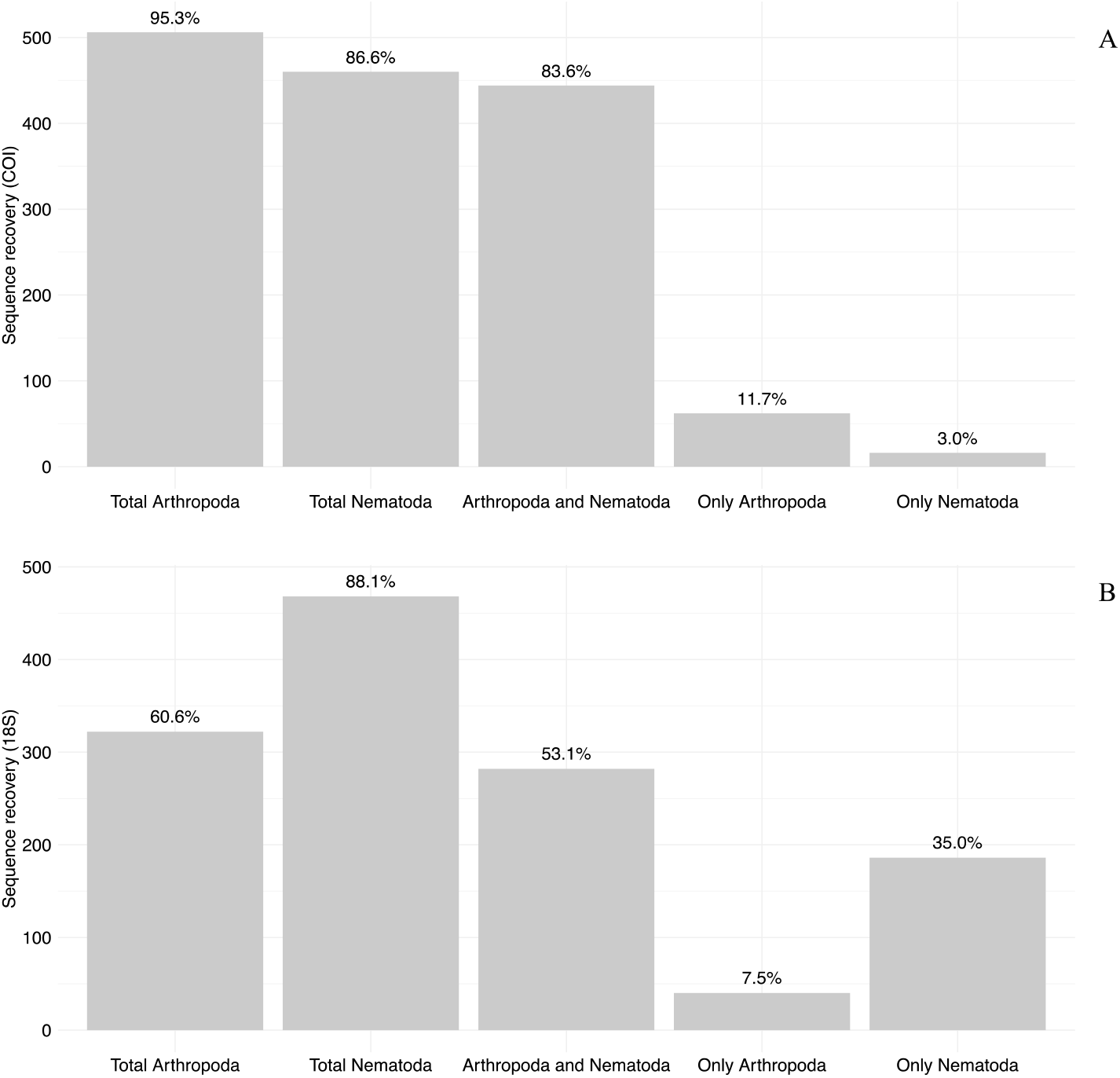
Histograms displaying sequence recovery for 531 arthropod specimens and associated nematodes assessed for COI (A) and 18S (B).

### Characterization of chimeras

Analysis of the downsized COI dataset with UCHIME identified 38 chimeric sequences, while 173 were identified in the 18S dataset (Supplementary table 2). All chimeras were unique, except for two COI chimeras that were each detected in two different samples: one in BIOUG88840-G11 and BIOUG89258-F01, and the other in BIOUG87363-H10 and BIOUG89258-F01. The number of chimeras characterized by a single crossover spot was 28 for COI and 144 for 18S, representing 73.7% and 83.2% of the total chimeras recovered with UCHIME. Analysis of the COI chimeric molecules with a single crossover site showed no evidence of crossover within the first or last ∼150 bp of the amplicon; they were restricted to the center of the sequence (Figure 2). The Kernel function revealed a high density of crossovers around 150–250 bp and a lower density around 450 bp from the 5’ end, but Hartigan’s Dip test did not reject the null hypothesis of unimodality (D = 0.11; p-value = 0.008) (Supplementary Figure 2). Similar to COI, analysis of crossover locations in the 144 chimeric 18S molecules characterized by a single crossover site showed no evidence of crossover in the first and last ∼200bp of the amplicon. Most crossovers were centrally located, approximately 300–550 bp from the 5’ end. However, this central region contained two distinct crossover hotspots: one located 300–400 base pairs, and another 500–550 base pairs from the 5’ end (Figure 2). Visual inspection of the Kernel density function confirmed these two distinct regions, revealing a bimodal distribution, further supported by Hartigan’s Dip test, which strongly rejected unimodality (Supplementary figure 2). ANOVA revealed a significant effect of parent sequence category (i.e., same phylum, different phyla, and unknown origin) on the width of crossover regions in 18S chimeras (df = 2, F = 20.57, p < 0.001) (Supplementary table 3). The taxonomic origins of the parent sequences display distinct patterns between COI and 18S markers. Twenty-six of 28 (92.9%) COI chimeras originated from sequences within the same phylum. The only two exceptions (7.1% of the total) included one chimera with an unknown origin, and another formed from a combination of arthropod and nematode sequences. Ninety-one of the 144 chimeric 18S (63.2%) sequences were formed from a host arthropod and an infecting nematode, 38 (26.4% of the total) originated from two sequences within the same phylum, and 15 (10.4% of the total) had an unknown origin.

**Figure 2.**
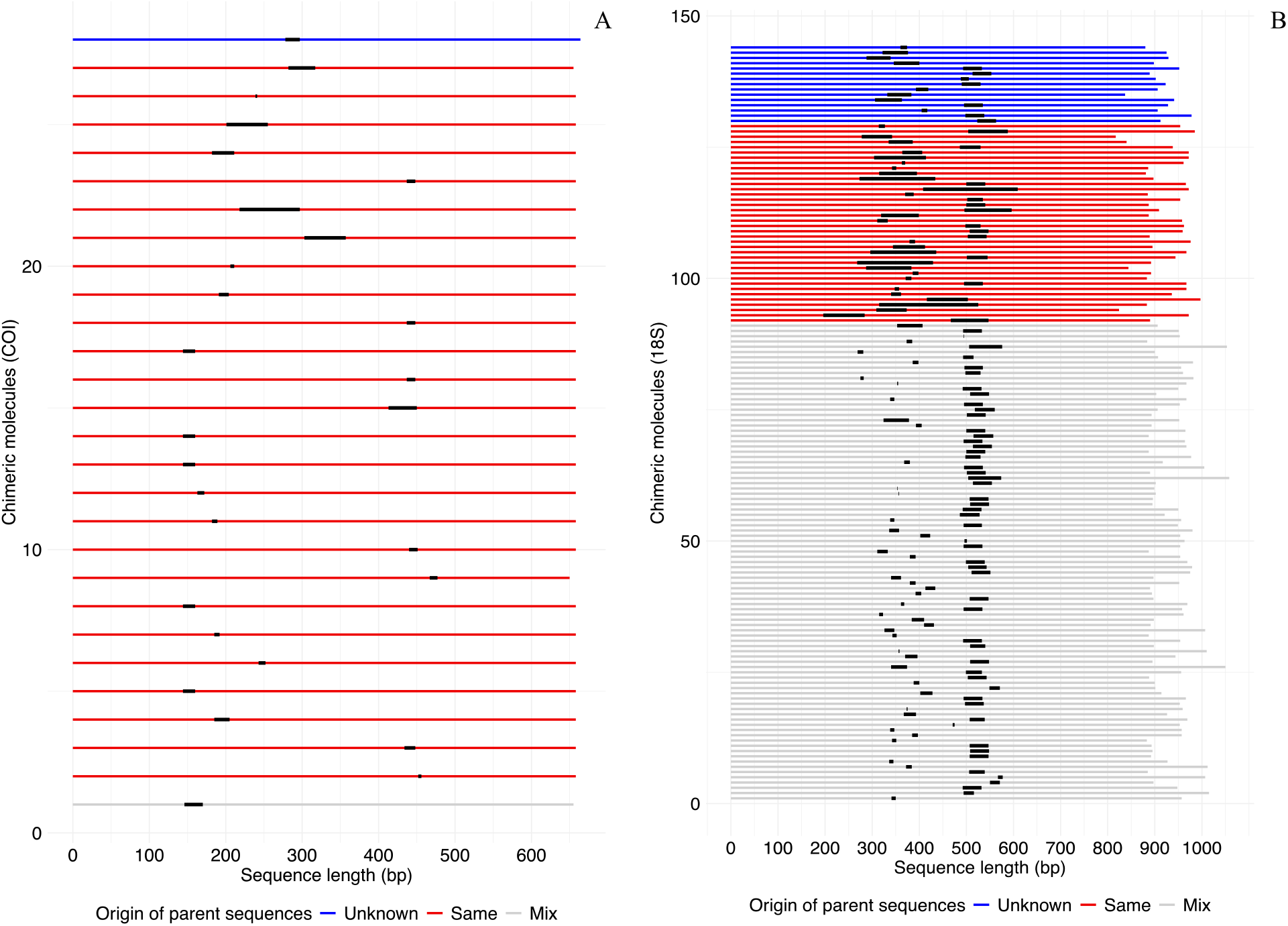
Location of the crossover ranges for the examined chimeric sequences for COI (A) and 18S (B). Each horizontal line represents a chimeric sequence, color-coded based on the classification of its parent sequences: Unknown (originating from unidentified phyla), Same (from the same phylum), Mix (from different phyla). Thick black lines indicate the crossover regions within each chimera.

### DNA stability and conserved regions

The average GC content of the consensus sequence derived from the aligned parent sequences of the COI chimeras was 23%, considerably lower than the 43% observed for 18S (Supplementary Figure 3). The consensus COI sequences had an average ΔG° of –1.1 kcal/mol with the lowest values reaching –1.3 kcal/mol (Figure 3). By comparison, the consensus of the 18S sequences had an average ΔG° of –1.32 kcal/mol, with two regions (300–400 bp & 500–550 bp from 5’ end) below –1.6 kcal/mol (Figure 3). GC content displayed an inverse relationship to DNA stability with regions of high GC content corresponding to areas of lower ΔG°, indicating increased stability (Supplementary figure 3).

**Figure 3.**
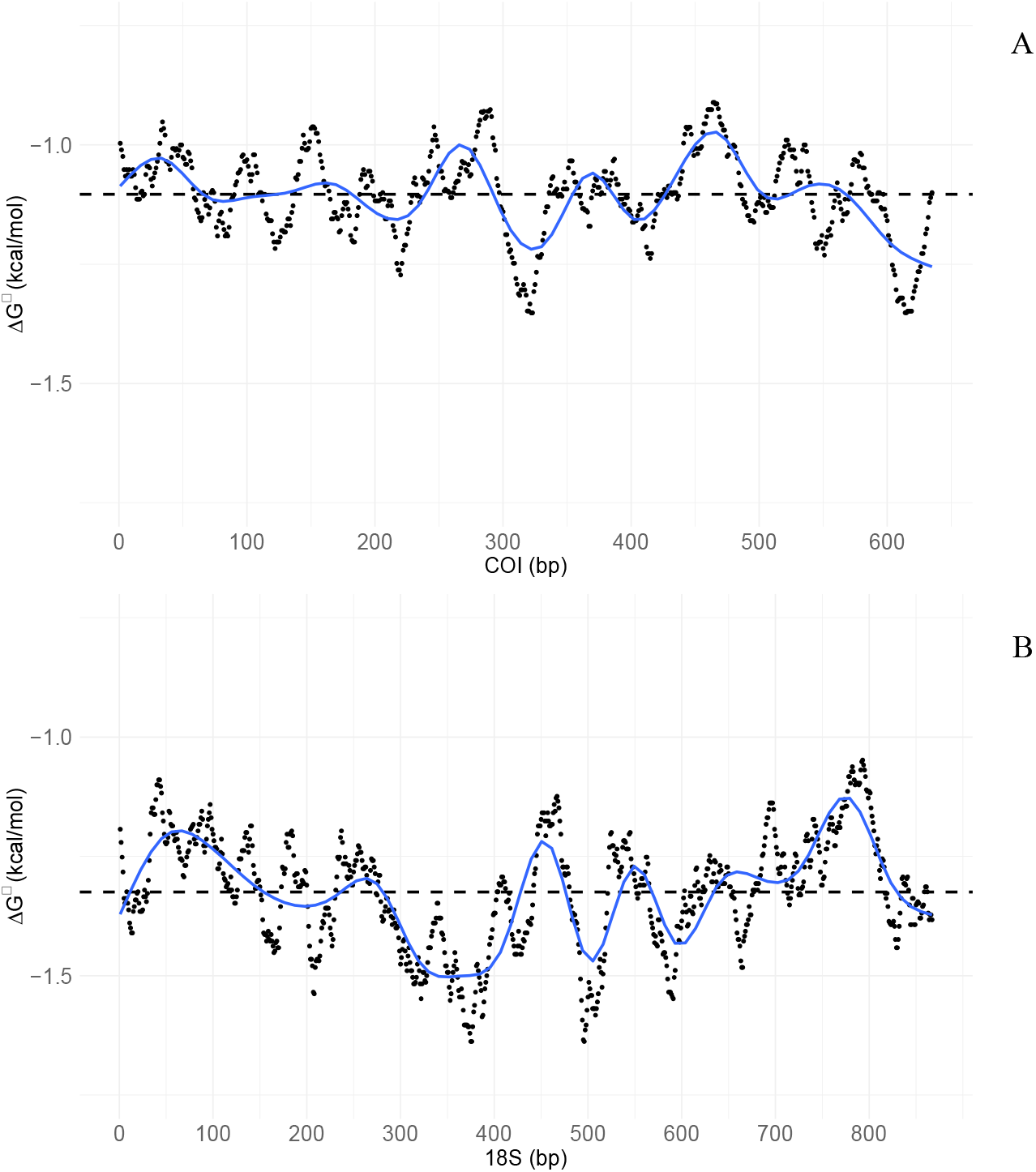
Sliding window analysis (window size 30 base pairs and step size 1 base pair) of DNA stability (measured as ΔG°) in the majority consensus sequences, COI (A) and 18S (B), derived from the alignments of parent sequences of the chimeras. Average value is represented by the dotted black line; the trendline is modelled using a GAM function.

The aligned parent sequences of the COI chimeras showed a fluctuating trend with a slight increase in identity (i.e., more conserved regions) at 150–250 bp, just after 400 bp, and around 600 bp from the 5’ end. By contrast, the aligned parent 18S sequences revealed two regions where identity frequently reaches 100% (i.e., perfectly conserved regions) at 300–400 bp and 500–550 bp from the 5’ end (Figure 4).

**Figure 4.**
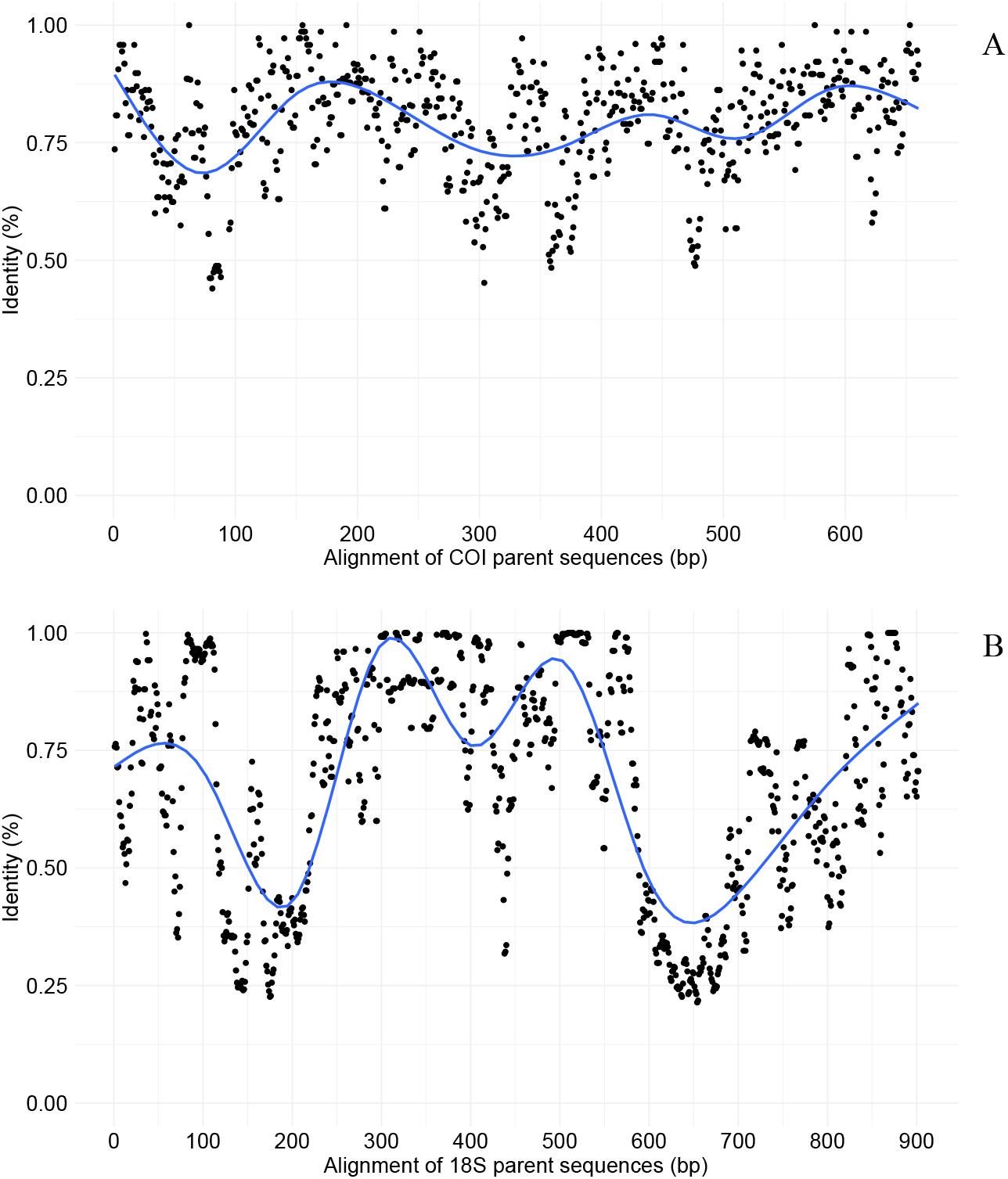
Sliding window analysis (window size 5 base pairs and step size 1 base pair) examining percentage of identity in the majority consensus sequences, COI (A) and 18S (B), derived from the alignments of parent sequences of the chimeras. The trendline is modelled using a GAM function.

### COI analysis

This analysis utilized the complete dataset of COI amplicons comprising total of 2,779,587 reads.

### Crossover regions

UCHIME identified 779 chimeric sequences of which 470 (60.3%) had a single crossover region. Crossover events were again absent in the first and last ∼150 bp of the chimeric amplicons, occurring exclusively in the central region. Closer inspection suggests a multimodal distribution, with a high density of crossovers near 200 bp that appears to comprise two closely spaced peaks. Another distinct region of increased density is observed around 450 bp. Hartigan’s dip test further supports this observation (D=0.058; p-value<0.01), indicating a departure from unimodality (Supplementary figure 4).

### DNA stability and conserved regions

The consensus sequence for the aligned COI sequences had an average GC content of 25% and an average ΔG° of –1.2 kcal/mol. Similar to the reduced COI and 18S datasets, GC content and ΔG° exhibited an inverse relationship. Notably, regions with high DNA stability, characterized by increased GC content (Supplementary figure 5) and lower ΔG° values (Figure 6), were observed near 200 bp, 300 bp, and 600 bp from the 5’ end.

**Figure 5.**
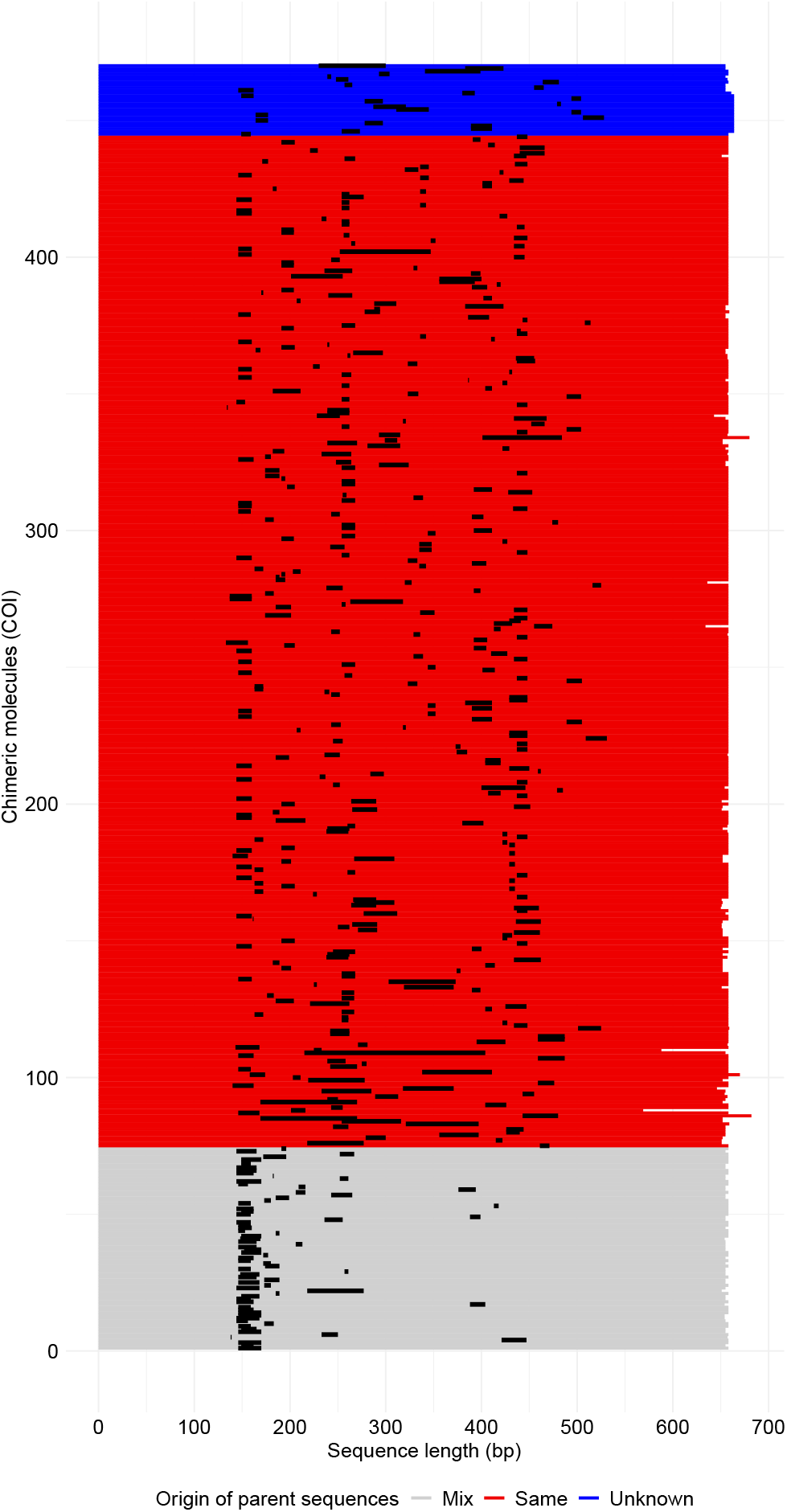
Location of the crossover ranges for the examined chimeric sequences for full COI dataset. Horizontal lines represent chimeric sequences which are color-coded to denote different categories of parent sequences; black thick lines represent crossover ranges.

**Figure 6.**
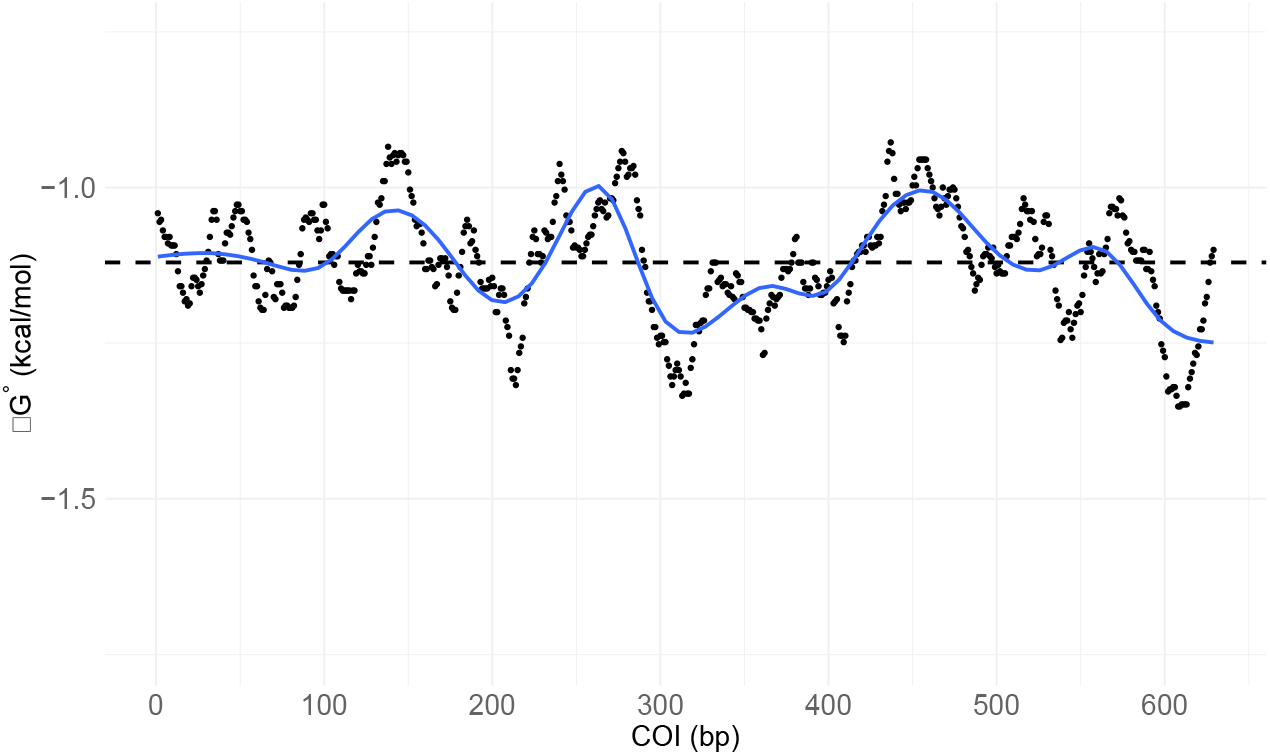
Sliding window analysis (window size 30 base pairs and step size 1 base pair) of DNA stability (measured as ΔG°) in the majority consensus sequence of the full dataset of COI, derived from the alignments of parent sequences of the chimeras. Average value is represented by the dotted black line; the trendline is modelled using a GAM function.

The alignment of parent sequences for the COI chimeras displayed a variable pattern, with higher sequence identity (i.e., more conserved regions) between 150–250 bp, just beyond 400 bp, and between 550–600 bp from the 5’ end (Figure 7).

**Figure 7.**
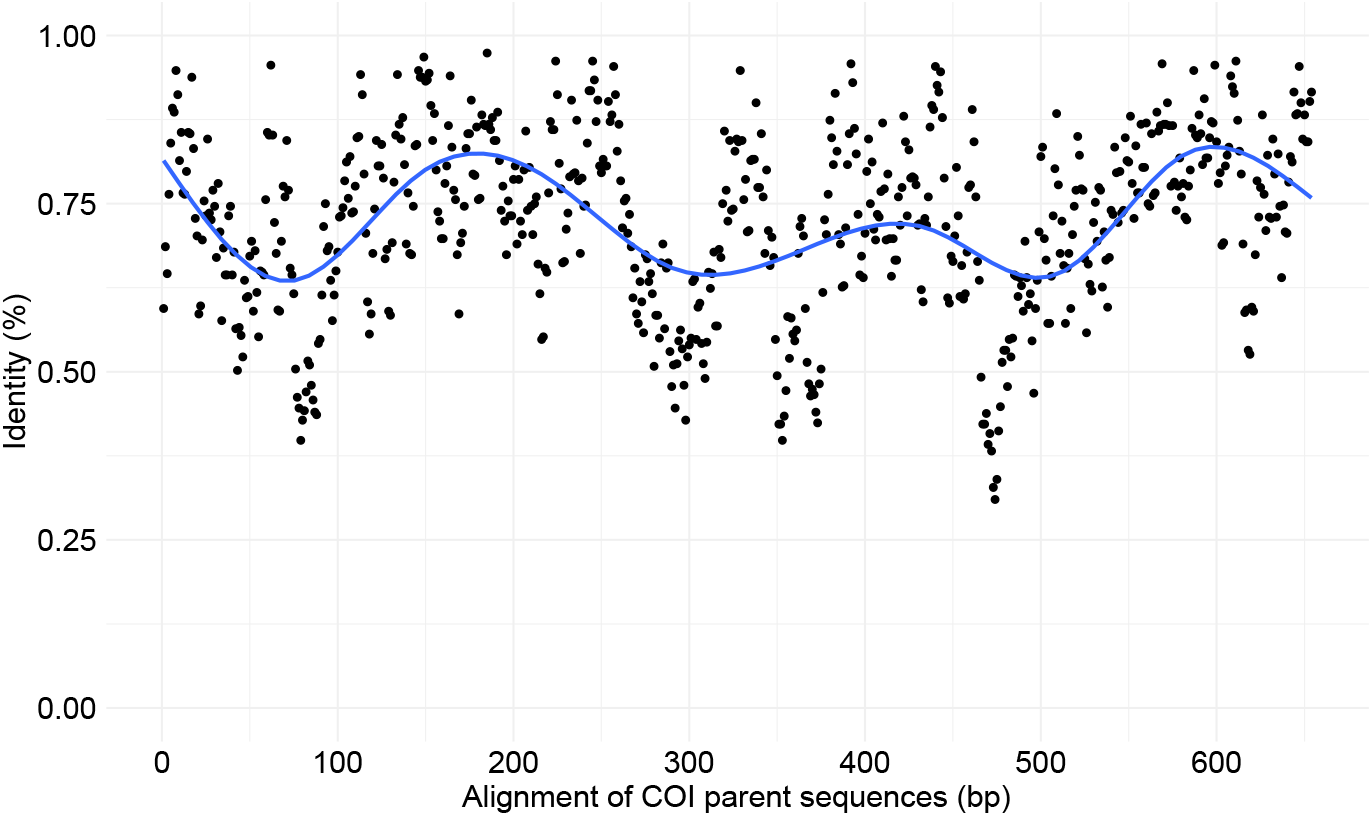
Sliding window analysis (window size 5 base pairs and step size 1 base pair) examining percentage of identity in the majority consensus sequence of the full COI dataset, derived from the alignments of parent sequences of the chimeras.

## Discussion

Sequencing of targeted gene regions is a widely adopted method for characterizing species diversity. However, accurate estimation of diversity can be impeded by methodological artefacts. This study examines the frequency and nature of chimeric sequences generated during PCR amplification of two genes (COI, 18S) in DNA extracts from arthropod specimens infected with nematodes.

The results revealed greater COI recovery from host arthropods (95.3%) than from their nematodes (86.6%). This pattern contrasts with the results for 18S, where recovery was higher for nematodes (88.1%) than their host arthropods (60.6%). The higher recovery of nematodes for 18S likely reflects the fact that the primers employed in this study were designed for nematodes (Floyd et al., 2002). The selection of the appropriate genetic markers and primer pairs is a critical aspect of eDNA/metabarcoding and symbiome studies. However, due to variation in target organisms, gene regions, PCR conditions, and sequencing platforms across studies, reaching general conclusions about the performance of different genetic markers is challenging. Comparisons between COI and 18S described in the literature reported distinct and even contradictory results. A study by Giebner et al. (2020), examining eDNA from freshwater samples and Malaise traps, revealed a similar performance between markers with a large overlap in recovered OTUs. By contrast, Leite et al. (2021) observed significant disparity in the species detected by each marker, with ∼70% of the species exclusively identified by either 18S or COI. In agreement with the present results, other studies have shown a taxon-dependent bias, with Arthropoda typically being preferentially recovered using COI and Nematoda having higher recovery with 18S (Atienza et al., 2020; Wangesteen et al., 2018; Sikder et al., 2020).

Chimeras are often prevalent (Qin et al., 2023), comprising up to 40% of PCR products in some studies (Porazinska et al., 2012; Lenz and Becker, 2008). The software employed for their detection splits sequences into smaller fragments, which are then queried against a reference dataset of chimera-free sequences (Haas et al., 2011; Edgar et al., 2016) or analyzed through a *de novo* approach (Wright et al., 2012; Mysara et al., 2015; Edgar et al., 2016). About a third of chimeras are undetected with these methods (Porazinska et al., 2012; Mysara et al., 2015), while approximately 2% are incorrectly identified as chimeras (Wright et al., 2012). This underscores the need for more effective detection methods and a clearer understanding of both the frequency of chimeras and the factors contributing to their formation. Comparison of the COI and 18S markers revealed 38 chimeric OTUs in COI and 174 in 18S. Given the much higher GC content of 18S (43%) than COI (23%), the elevated incidence of 18S chimeras confirms that high GC content facilitates chimera formation (Qin et al., 2023). Despite this association, the underlying reasons for this association has never been examined. Our results suggest that two interconnected mechanisms influence chimera formation. First, the prerequisite for the genesis of chimeras is the formation of incompletely extended strands. While Taq DNA polymerase typically adds about 50 nucleotides per binding event (Merkens et al., 1995), regions of high DNA stability can increase the likelihood of polymerase dissociation. This study supports this hypothesis by demonstrating the correspondence between regions of high DNA stability (i.e., low ΔG°) and crossover hotspots, with the18S gene revealing two distinct regions between 300–400 base pairs and 500–550 base pairs from the 5’ end. Second, chimera formation requires that incompletely extended strands bind to a different template. Conserved regions play a crucial role in this, as the 3’ end of the incomplete strand must bind to template from another species and continue extension. This study revealed a correspondence between conserved regions and crossover hotspots, with 18S once again displaying distinct regions between 300–400 base pairs and 500–550 base pairs from the 5’ end. Although the reduced COI dataset showed a pattern similar to that for 18S, establishing a clear correlation between crossover hotspots, DNA stability, and conserved regions was challenging because so few chimeras were detected (28). While the full COI dataset did not reveal a correlation between DNA stability and crossover regions, it did show a correlation between conserved regions and crossover hotspots. For COI, a broad peak near 200 bp appears to be comprised of two close sub-peaks, along with a distinct peak around 400 bp. This result suggests that sequence identity has a greater influence than DNA stability, particularly for genes, like COI, with low DNA stability and low GC content throughout the sequence. Notably, no chimeras were detected near the termini of COI and 18S, possibly due to the steady extension during the first 150 bp of the template. 18S chimeras also revealed that the taxonomy of the parent sequences has a highly significant effect on the width of the crossover range for 18S (F=20.57; p-value<0.01). This is expected, as the lower sequence divergence within arthropod and within nematode lineages makes it difficult to pinpoint the exact crossover location. In contrast, the high genetic divergence between these two distant phyla facilitates precise identification of crossover points.

The formation of chimeras during PCR involves the concatenation of different template sequences, creating sequences not present in the original DNA extract. These artefactual sequences raise estimates of total projected diversity because both specialized parametric models (Zhang et al., 2010) and general non-parametric approaches (Phillips et al., 2020; Hsieh et al., 2016) rely on observed diversity to estimate total diversity (D’Ercole et al., 2021; Dincă et al., 2021). In the present study, the oversight of chimeras would have inflated COI diversity by 3.6% and 18S by 16.3%. It needs emphasis that this study primarily focused on chimeras derived from organisms in different phyla. However, chimeras are both more frequent and much harder to detect when closely related species are involved (Haas et al., 2011; Wright et al., 2012). Detection gains further complexity in organisms with many NUMTs (Hebert et al., 2023; Schultz et al., 2022) or paralog genes (Kaltenegger and Ober, 2015) as chimeras can easily form between these intra-individual sequence variants. The present study deepens understanding of both the frequency of chimeras and the factors contributing to their formation, providing insights that can inform and improve detection algorithms. The identified crossover hotspots for COI and 18S can be used to specifically examine the flanking DNA for more accurate detection. Additionally, this study highlights key sequence features, such as DNA stability and highly conserved regions, which aid prediction of crossover regions.

## Supporting information

Supplementary script 1

Supplementary script 2

Supplementary script 3

Supplementary script 4

## Acknowledgments

This study was supported by the Government of Canada through Genome Canada and Ontario Genomics (OGI-208, OGI-233), the New Frontiers in Research Fund (NFRFT-2020-00073), the Canada Foundation for Innovation (MSI), and by the Ontario Ministry of Colleges and Universities.

We also thank Andreas Kolter for his suggestions on earlier versions of the manuscript.

## Supplementary materials

**Supplementary figure 1**. Histograms displaying the number of OTUs per sample detected for COI (A) and 18S (B).

**Supplementary figure 2**. Kernel density estimate of the mid-point values of the crossover ranges of chimeras for COI (A) and 18S (B). Dashed lines indicate the mean value.

**Supplementary figure 3**. Sliding window analysis (window size 30 base pairs and step size 1 base pair) of GC content in the majority consensus sequences, COI (A) and 18S (B), derived from the alignments of parent sequences of the chimeras. Average value is represented by the dotted black line; the trendline is modelled using a GAM function.

**Supplementary figure 4**. Kernel density estimate of the mid-point values of the crossover ranges of chimeras for the full COI dataset.

**Supplementary figure 5**. Sliding window analysis (window size 30 base pairs and step size 1 base pair) of GC content in the majority consensus sequence of the full COI dataset, derived from the alignments of parent sequences of the chimeras. Average value is represented by the dotted black line; the trendline is modelled using a GAM function.

**Supplementary table 1**. List of samples with corresponding PAD sequences, UMIs (Unique Molecular Identifiers), and primer sequences, for both COI and 18S.

**Supplementary table 2**. List of samples with corresponding COI and 18S sequences, including taxonomic classification, sequence read counts, and sequence lengths. Rows representing chimeric sequences identified by UCHIME are highlighted in orange.

**Supplementary table 3**. Analysis of variance (ANOVA) examining differences in metrics such as mid-point of crossover regions and width of crossover ranges among different categories such as host organism and source of the parent sequences.

**Supplementary script 1**. R script used to display the crossover range of examined chimeric sequences and to color-code them based on the origin of their parental sequences.

**Supplementary script 2**. R script performing a sliding window analysis (30 bp window size, 1 bp step size) of GC content to estimate the percentage of sequence identity within a single alignment.

**Supplementary script 3**. R script that performs a sliding window analysis (30 bp window size, 1 bp step size) to estimate DNA stability, measured as ΔG°, in the majority consensus sequences of the parental molecules of chimeras.

**Supplementary script 4**. R script performing a sliding window analysis (5 bp window size, 1 bp step size) to estimate percentage of identity in the majority consensus sequences of the parental molecules of chimeras.

## References

Acinas SG, Klepac-Ceraj V, Hunt DE, Pharino C, Ceraj I, Distel DL, Polz MF. 2004. Fine-scale phylogenetic architecture of a complex bacterial community. Nature. 430(6999):551–554.

Ashelford KE, Chuzhanova NA, Fry JC, Jones AJ, Weightman AJ. 2006. New screening software shows that most recent large 16S rRNA gene clone libraries contain chimeras. Applied and Environmental Microbiology. 72(9):5734–5741.

Atienza S, Guardiola M, Præbel K, Antich A, Turon X, Wangensteen OS. 2020. DNA metabarcoding of deep-sea sediment communities using COI: community assessment, spatiotemporal patterns and comparison with 18S rDNA. Diversity. 12(4):123.

Bass D, Rueckert S, Stern R, Cleary AC, Taylor JD, Ward GM, Huys R. 2021. Parasites, pathogens, and other symbionts of copepods. Trends in Parasitology. 37(10):875–889.

Burns JM, Janzen DH, Hajibabaei M, Hallwachs W, Hebert PDN. 2008. DNA barcodes and cryptic species of skipper butterflies in the genus Perichares in Area de Conservacion Guanacaste, Costa Rica. Proceedings of the National Academy of Sciences USA. 105(17):6350–6355.

Cruaud P, Rasplus JY, Rodriguez LJ, Cruaud A. 2017. High-throughput sequencing of multiple amplicons for barcoding and integrative taxonomy. Scientific Reports. 7(1):41948.

D’Ercole J, Dincă V, Opler PA, Kondla NG, Schmidt C, Phillips JD, Robbins R, Burns JM, Miller SE, Grishin NV, Zakharov EV, deWaard JR, Ratnasingham S, Hebert PDN. 2021. A DNA barcode library for the butterflies of North America. PeerJ. 9:e11157.

D’Ercole J, Dapporto L, Schmidt C, Dincă V, Talavera G, Vila R, Hebert PDN. 2022. Patterns of DNA barcode diversity in butterfly species (Lepidoptera) introduced to the Nearctic. European Journal of Entomology. 119:379–87.

D’Ercole J, Dapporto L, Opler P, Schmidt CB, Ho C, Menchetti M, Zakharov EV, Burns JM, Hebert PDN. 2024. A genetic atlas for the butterflies of continental Canada and United States. PLoS ONE. 19(4): e0300811.

Dapporto L, Menchetti M, Vodă R, Corbella C, Cuvelier S, Djemadi I, Gascoigne-Pees M, Hinojosa JC, Lam NT, Serracanta M, Talavera G, Dincă V, Vila R. 2022. The atlas of mitochondrial genetic diversity for Western Palaearctic butterflies. Global Ecology and Biogeography. 31(11):2184–2190.

Dapporto L, Menchetti M, Dincă V, Talavera G, Garcia-Berro A, D’Ercole J, Hebert PD, Vila R. 2024. The genetic legacy of the Quaternary ice ages for West Palearctic butterflies. Science Advances. 10(38): eadm8596.

Dinca V, Dapporto L, Somervuo P, Vodă R, Cuvelier S, Gascoigne-Pees M, Huemer P, Mutanen M, Hebert PDN, Vila R. 2021. High resolution DNA barcode library for European butterflies reveals continental patterns of mitochondrial genetic diversity. Communications Biology. 4:315.

Edgar RC, Haas BJ, Clemente JC, Quince C, Knight R. 2011. UCHIME improves sensitivity and speed of chimera detection. Bioinformatics. 27(16):2194–2200.

Edgar RC. 2016. UCHIME2: improved chimera prediction for amplicon sequencing. BioRxiv. DOI: 10.1101/074252.

Floyd R, Abebe E, Papert A, Blaxter M. 2002. Molecular barcodes for soil nematode identification. Molecular Ecology. 11(4):839–580.

Giebner H, Langen K, Bourlat SJ, Kukowka S, Mayer C, Astrin JJ, Misof B, Fonseca VG. 2020. Comparing diversity levels in environmental samples: DNA sequence capture and metabarcoding approaches using 18S and COI genes. Molecular Ecology Resources. 20(5):1333–1345.

Haas BJ, Gevers D, Earl AM, Feldgarden M, Ward DV, Giannoukos G, Ciulla D, Tabbaa D, Highlander SK, Sodergren E, Methé B, DeSantis TZ, Human Microbiome Consortium, Petrosino JF, Knight R, Birren BW. 2011. Chimeric 16S rRNA sequence formation and detection in Sanger and 454-pyrosequenced PCR amplicons. Genome Research. 21(3):494–504.

Hardulak LA, Morinière J, Hausmann A, Hendrich L, Schmidt S, Doczkal D, Müller J, Hebert PDN, Haszprunar G. 2020. DNA metabarcoding for biodiversity monitoring in a national park: screening for invasive and pest species. Molecular Ecology Resources. 20:1542–1557.

Hartigan JA, Hartigan PM. 1985. The dip test of unimodality. The Annals of Statistics. 1:70–84.

Hebert PDN, Cywinska A, Ball SL, deWaard JR. 2003. Biological identifications through DNA barcodes. Proceedings of the Royal Society B: Biological Sciences. 270(1512):313–321.

Hebert PDN, Penton EH, Burns JM, Janzen DH, Hallwachs W. 2004. Ten species in one: DNA barcoding reveals cryptic species in the neotropical skipper butterfly Astraptes fulgerator. Proceedings of the National Academy of Sciences USA. 101(41):14812–14817.

Hebert PDN, Braukmann TW, Prosser SWJ, Ratnasingham S, DeWaard JR, Ivanova NV, Janzen DH, Hallwachs W, Naik S, Sones JE, Zakharov EV. 2018. A Sequel to Sanger: amplicon sequencing that scales. BMC Genomics. 19:1–14.

Hebert PDN, Bock DG, Prosser SWJ. 2023. Interrogating 1000 insect genomes for NUMTs: A risk assessment for estimates of species richness. PLoS One. 18(6):e0286620.

Hebert PDN, Floyd R, Jafarpour S, Prosser SWJ. 2024. Barcode 100K specimens: In a single Nanopore run. Molecular Ecology Resources. 30:e14028.

Hsieh TC, Ma KH, Chao A. 2016. iNEXT: an R package for rarefaction and extrapolation of species diversity (Hill numbers). Methods in Ecology and Evolution. 7:1451–1456.

Kalle E, Kubista M, Rensing C. 2014. Multi-template polymerase chain reaction. Biomolecular Detection and Quantification. 2:11–29.

Kaltenegger E, Ober D. 2015. Paralogue interference affects the dynamics after gene duplication. Trends in Plant Science. 20(12):814–821.

Katoh K, Misawa K, Kuma K, Miyata T. 2002. MAFFT: a novel method for rapid multiple sequence alignment based on fast Fourier transform. Nucleic Acids Research. 30(14):3059–3066.

Leite BR, Vieira PE, Troncoso JS, Costa FO. 2021. Comparing species detection success between molecular markers in DNA metabarcoding of coastal macroinvertebrates. Metabarcoding and Metagenomics. 5:249–260.

Lenz TL, Becker S. 2008. Simple approach to reduce PCR artefact formation leads to reliable genotyping of MHC and other highly polymorphic loci—implications for evolutionary analysis. Gene. 427(1-2):117–123.

Liu J, Jiang J, Song S, Tornabene L, Chabarria R, Naylor GJ, Li C. 2017. Multilocus DNA barcoding–species identification with multilocus data. Scientific Reports. 7(1):16601.

Merkens LS, Bryan SK, Moses RE. 1995. Inactivation of the 5’-3’ exonuclease of Thermus aquaticus DNA polymerase. Biochimica et Biophysica Acta (BBA)—Gene Structure and Expression. 1264(2):243–248.

Minardi D, Ryder D, Del Campo J, Garcia Fonseca V, Kerr R, Mortensen S, Pallavicini A, Bass D. 2022. Improved high throughput protocol for targeting eukaryotic symbionts in metazoan and eDNA samples. Molecular Ecology Resources. 22(2):664–768.

Mysara M, Saeys Y, Leys N, Raes J, Monsieurs P. 2015. CATCh, an ensemble classifier for chimera detection in 16S rRNA sequencing studies. Applied and Environmental Microbiology. 81(5):1573–1584.

Phillips JD, French SH, Hanner RH, Gillis DJ. 2020. HACSim: an R package to estimate intraspecific sample sizes for genetic diversity assessment using haplotype accumulation curves. PeerJ Computer Science 6:e243.

Porazinska DL, Giblin-Davis RM, Sung W, Thomas WK. 2012. The nature and frequency of chimeras in eukaryotic metagenetic samples. The Journal of Nematology. 44(1):18–25.

Prosser SW, Floyd RM, Thompson KA, Hebert PDN. 2025. BOLDistilled: Comprehensive but compact DNA barcode reference libraries. EcoEvoRxiv.

Qin Y, Wu L, Zhang Q, Wen C, Van Nostrand JD, Ning D, Raskin L, Pinto A, Zhou J. 2023. Effects of error, chimera, bias, and GC content on the accuracy of amplicon sequencing. Msystems. 8(6):e0102523.

Robertson JM, Walsh-Weller J. 1998. An introduction to PCR primer design and optimization of amplification reactions. Methods in Molecular Biology. 98:121–154.

Rodríguez-Ezpeleta N, Zinger L, Kinziger A, Bik HM, Bonin A, Coissac E, Emerson BC, Martins Lopes C, Pelletier TA, Taberlet P, Narum S. 2021. Biodiversity monitoring using environmental DNA. Molecular Ecology Resources. 21(5):1405–1409.

Rognes TT, Flouri B, Nichols C, Quince Mahé F. 2016. VSEARCH: a versatile open source tool for metagenomics. PeerJ. 4:e2584.

Santa Lucia Jr J. 1998. A unified view of polymer, dumbbell, and oligonucleotide DNA nearestneighbor thermodynamics. Proceedings of the National Academy of Sciences USA. 95(4):1460– 1465.

Schultz JA, Hebert PDN. 2022. Do pseudogenes pose a problem for metabarcoding marine animal communities? Molecular Ecology Resources. 22(8):2897–2914.

Sikder MM, Vestergård M, Sapkota R, Kyndt T, Nicolaisen M. 2020. Evaluation of metabarcoding primers for analysis of soil nematode communities. Diversity. 12(10):388.

Thompson JR, Marcelino LA, Polz MF. 2002. Heteroduplexes in mixed-template amplifications: formation, consequence and elimination by ‘reconditioning PCR’. Nucleic Acids Research. 30(9):2083–2088.

Van Der Heyde M, Bunce M, Wardell-Johnson G, Fernandes K, White NE, Nevill P. 2020. Testing multiple substrates for terrestrial biodiversity monitoring using environmental DNA metabarcoding. Molecular Ecology Resources. 20(3):732–745.

Wang GCY, Wang Y. 1996. The frequency of chimeric molecules as a consequence of PCR coamplification of 16S rRNA genes from different bacterial species. Microbiology. 142:1107–1114.

Wangensteen OS, Palacín C, Guardiola M, Turon X. 2018. DNA metabarcoding of littoral hardbottom communities: high diversity and database gaps revealed by two molecular markers. PeerJ. 6:e4705.

Wickham H. 2016. ggplot2: elegant graphics for data analysis. Springer-Verlag: New York.

Wood SN. 2011. Fast stable restricted maximum likelihood and marginal likelihood estimation of semiparametric generalized linear models. Journal of the Royal Statistical Society (B). 73(1): 3– 36.

Wright ES, Yilmaz LS, Noguera DR. 2012. DECIPHER, a search-based approach to chimera identification for 16S rRNA sequences. Applied and Environmental Microbiology. 78(3):717–725.

Zhang AB, He LJ, Crozier RH, Muster C, Zhu CD. 2010. Estimating sample sizes for DNA barcoding. Molecular Phylogenetics and Evolution. 54(3):1035–1039.

