## Supplementary figures and images for "Incidence and attributes of chimeric COI and 18S sequences derived from nematode-infected arthropods"

### Supplementary figure 1

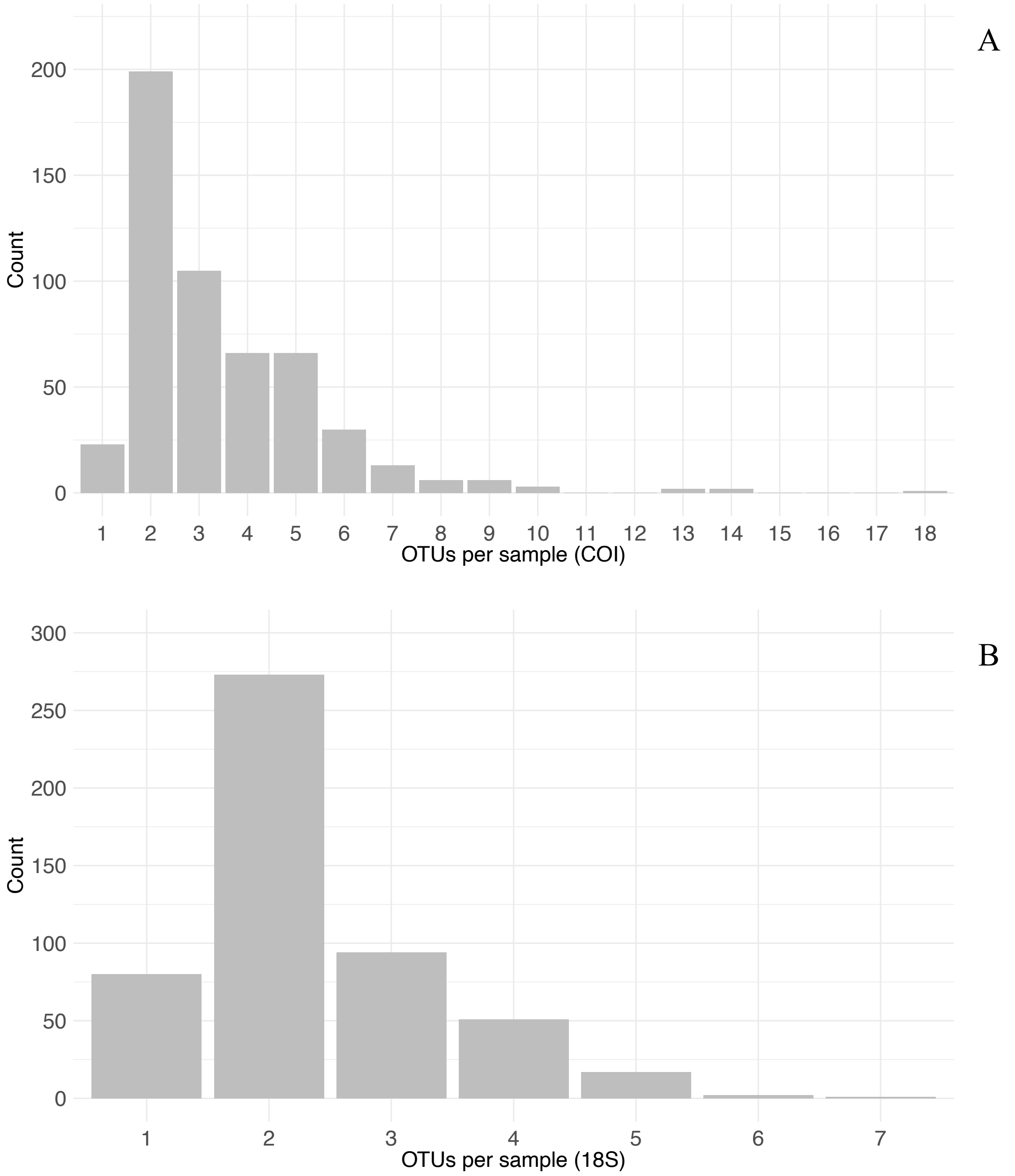

### Supplementary figure 2

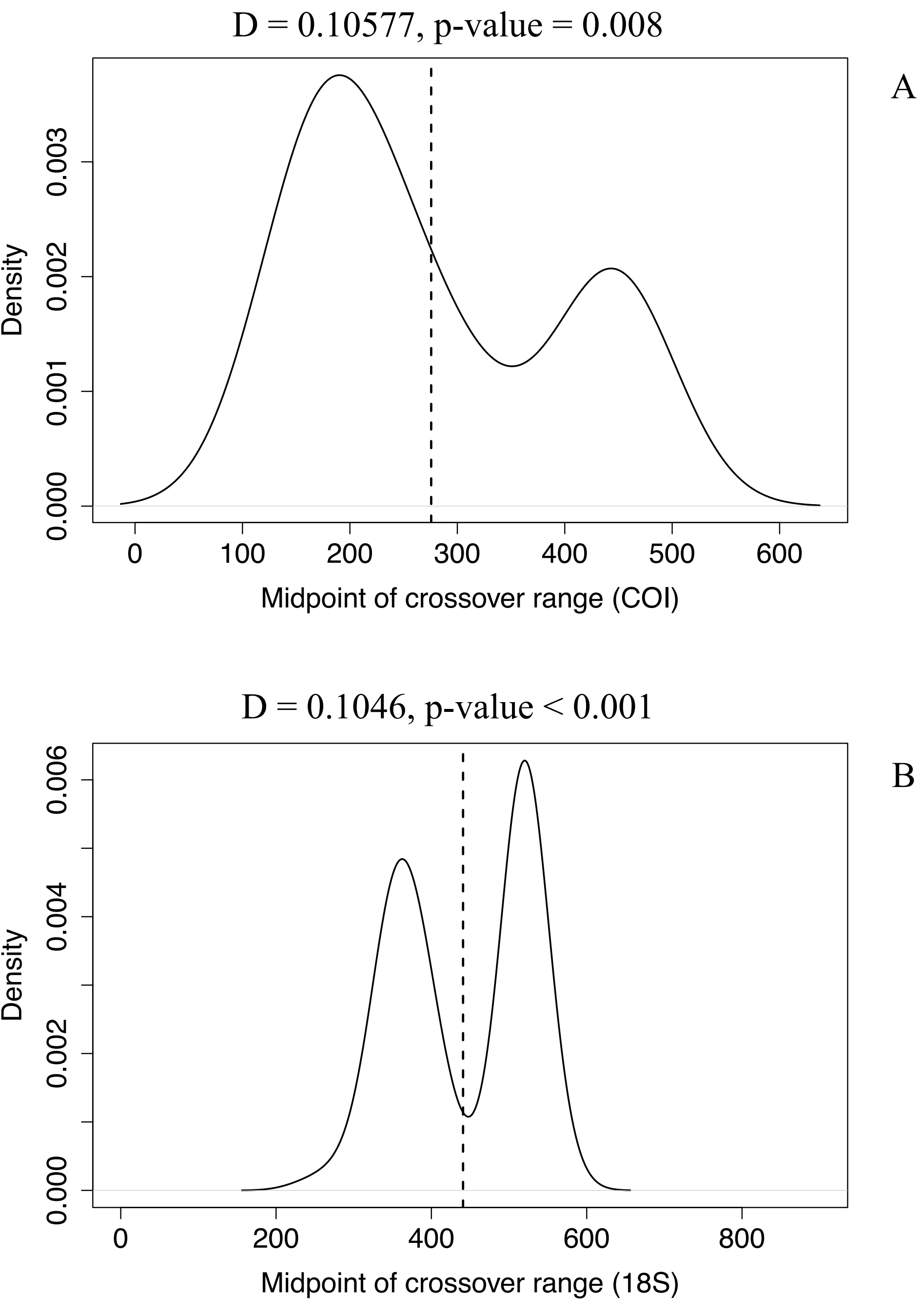

### Supplementary figure 3

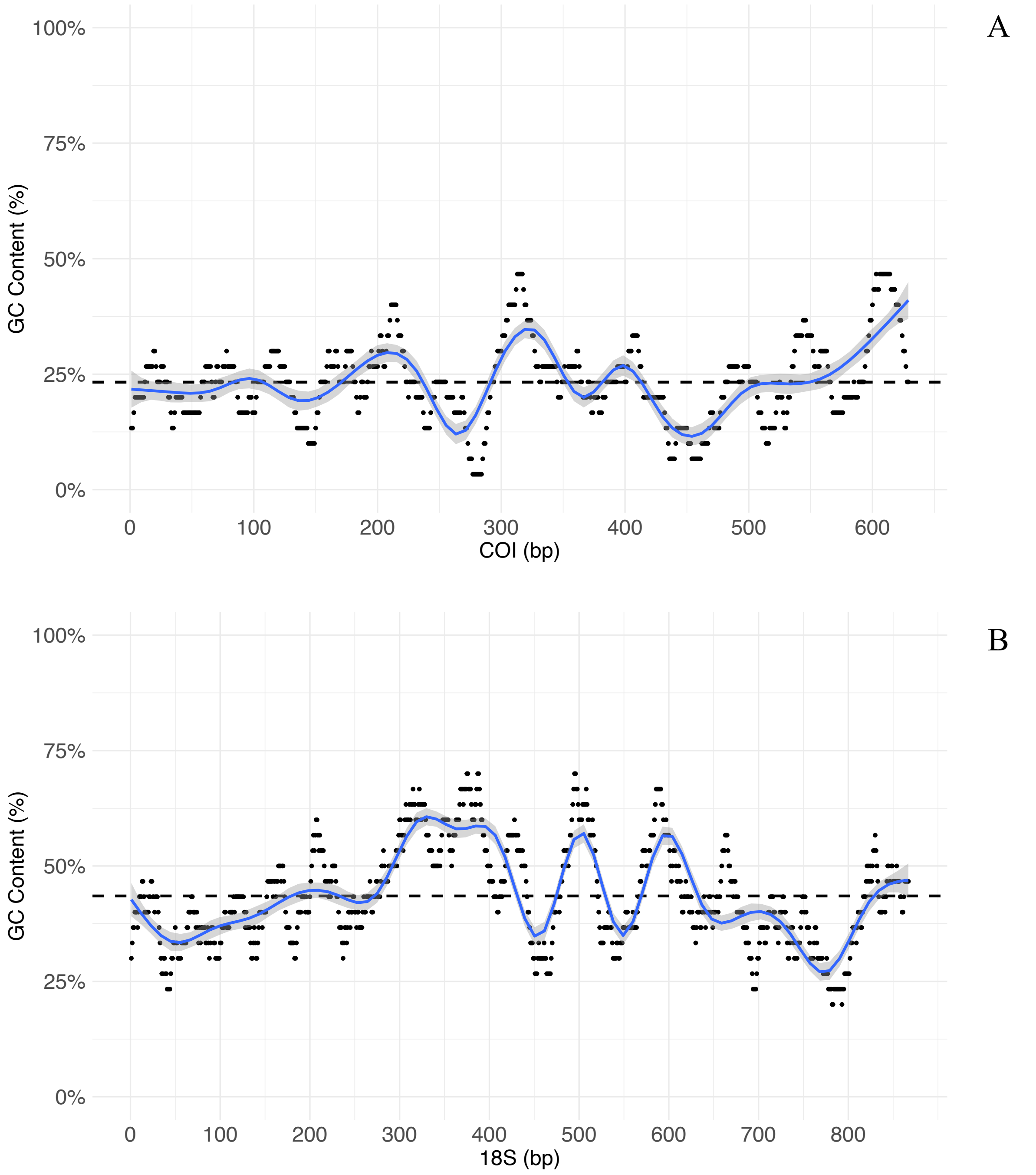

### Supplementary figure 4

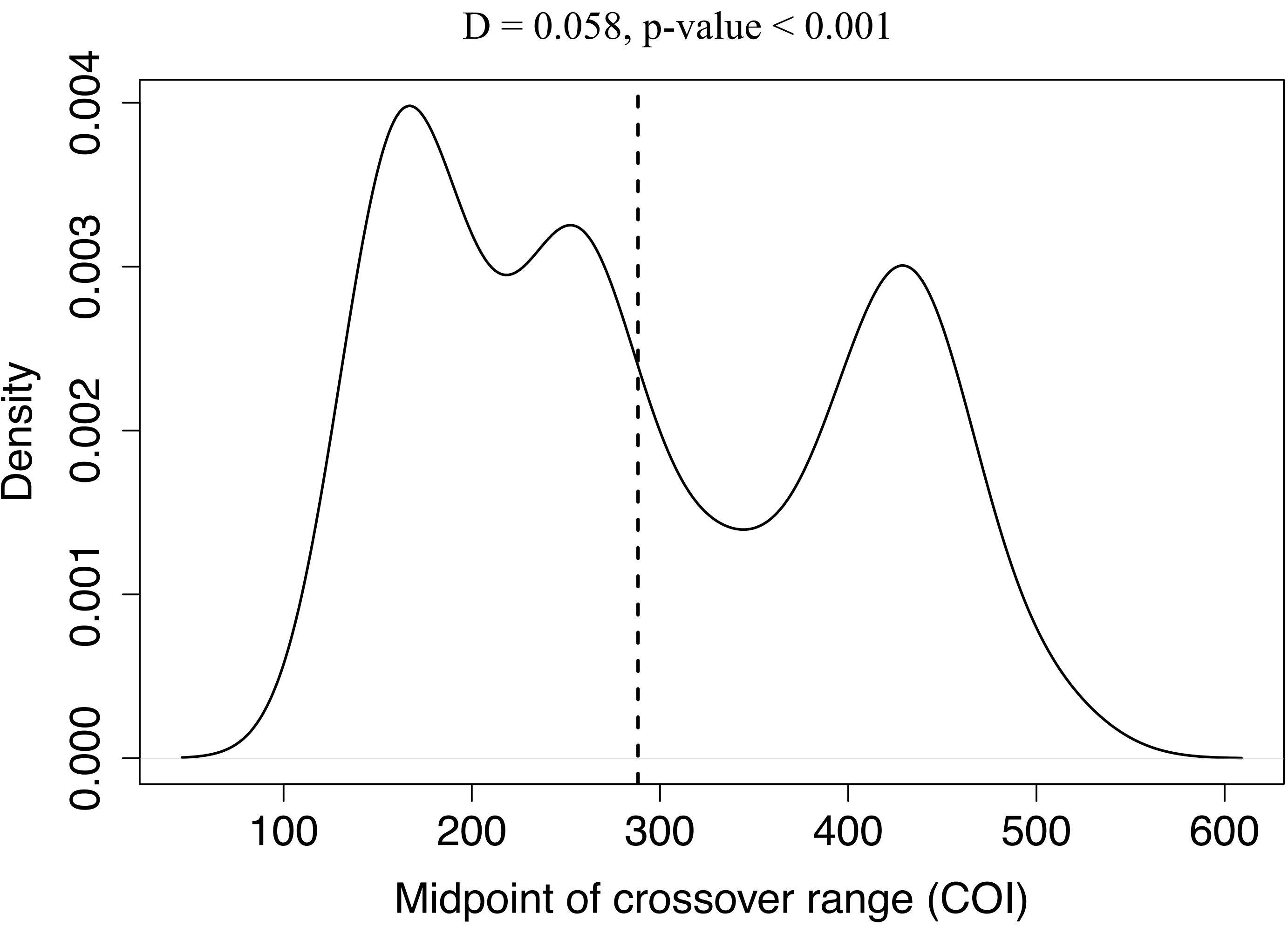

### Supplementary figure 5

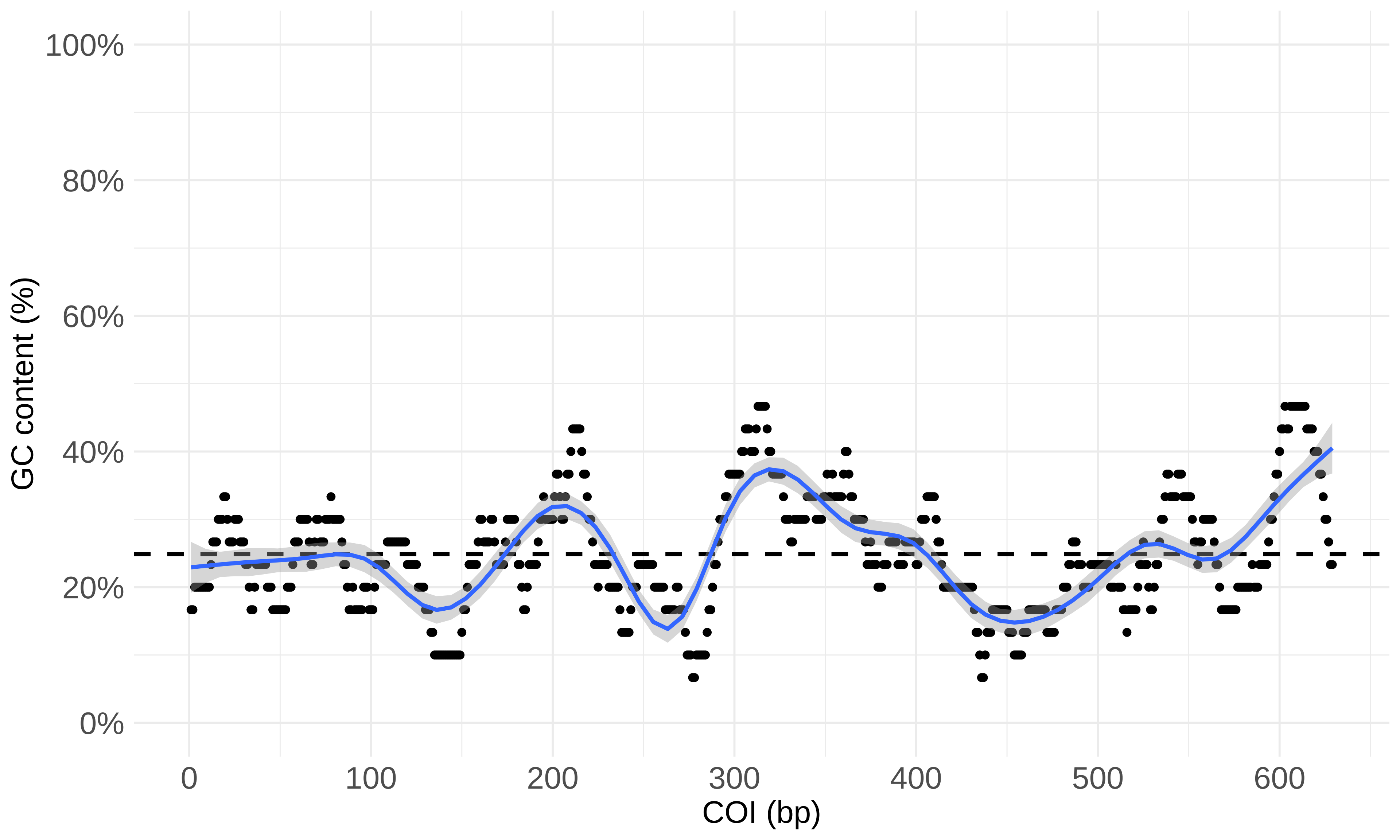
