## Supplementary script 1 for "Incidence and attributes of chimeric COI and 18S sequences derived from nematode-infected arthropods"

#Graphical representation of crossover ranges and parental origin of chimeric sequences

\# Required libraries

library(readxl)

library(ggplot2)

\# Input file description:

\# The Excel file should contain the following columns:

\# - 'recomb_range': a string of the form "start-end" (e.g., "145-202")

\# - 'seq_length': total length of the chimeric molecule in base pairs

\# - 'system': one of "Unknown", "Same", or "Mix", describing the origin of the parent sequences

\# Step 1: Read Excel file

file_path <- file.choose() # Prompt user to select the input Excel file

dataset <- read_excel(file_path)

\# Step 2: Extract recombination breakpoints

start_end <- sapply(dataset$recomb_range, function(x) {

as.numeric(unlist(strsplit(x, "-")))

})

dataset$start <- start_end[1, ]

dataset$end <- start_end[2, ]

\# Step 3: Set plotting order of systems

dataset$system <- factor(dataset$system, levels = c("Unknown", "Same", "Mix"))

\# Sort so "Unknown" appears at the top, followed by "Same", then "Mix"

dataset <- dataset[order(dataset$system, decreasing = TRUE), ]

\# Step 4: Define plot colors

colors <- c("Mix" = "grey77", "Same" = "red2", "Unknown" = "blue")

\# Step 5: Create the plot

plot <- ggplot(dataset, aes(y = seq_along(seq_length), x = seq_length, color = system)) +

geom_segment(aes(yend = seq_along(seq_length), x = 0, xend = seq_length), size = 1) + # Full sequence line

geom_segment(aes(x = start, xend = end, yend = seq_along(seq_length)), color = "black", size = 1.5) + # Recombination region

scale_color_manual(values = colors, drop = FALSE) +

scale_x_continuous(breaks = seq(0, 1050, by = 100)) +

labs(

y = "Chimeric molecules (COI)",

x = "Sequence length (bp)",

color = "Origin of parent sequences"

) +

theme_minimal() +

theme(

legend.position = "bottom",

axis.title = element_text(size = 16, color = "black"),

axis.text = element_text(size = 16, color = "black"),

legend.title = element_text(size = 16, color = "black"),

legend.text = element_text(size = 16, color = "black"),

plot.margin = margin(20, 20, 20, 20)

)

\# Step 6: Display the plot

print(plot)
