## Supplementary script 2 for "Incidence and attributes of chimeric COI and 18S sequences derived from nematode-infected arthropods"

\# Sliding Window GC Content Analysis

\# Description: Calculates and plots GC content across a DNA sequence using a sliding window approach.

\# Load required packages

library(Biostrings)

library(ggplot2)

library(writexl)

library(gam)

\# ----- Define genetic marker -----

marker_name <- "COI"

cat("Marker set to:", marker_name, "\n")

\# ----- Define DNA sequence -----

\# Replace this string with your sequence of interest

dna_seq <- DNAString("AACTTTATATTTTATTTTTGGTATTTGATCAGGAATAATTGGATCATCTTTAAGAATATTAATTCGATTAGAATTAAGAAAACCTGGTTCTTTAATTGGTAATGATCAAATTTATAATGTAATTGTTACAGCTCATGCTTTTATTATAATTTTTTTTATAGTTATACCTATTATAATTGGAGGATTTGGAAATTGATTAATTCCTTTAATATTAGGAGCTCCTGATATAGCTTTTCCTCGAATAAATAATATAAGATTTTGATTATTACCTCCTTCATTAATTTTATTATTATTTAGAAGTTTAATTGATAATGGAGCAGGAACAGGATGAACTGTTTATCCTCCTTTATCTTCTAATATTGCTCATAGTGGTACTTCAGTTGATTTAGCTATTTTTTCTTTACATTTAGCAGGAATTTCTTCAATTTTAGGAGCAATTAATTTTATTTCTACAATTATTAATATACGAACTAATAATATTAAATTAGATCAAATTCCTTTATTTGTTTGATCAGTTTTAATTACAGCTATTTTATTACTTTTATCTTTACCAGTTTTAGCTGGAGCTATTACTATATTATTAACTGATCGAAATTTAAATACTTCATTTTTTGATCCTGCAGGAGGAGGAGATCCAATTTTATATCAACATTTATTT")

\# ----- Sliding window parameters -----

window_size <- 30 # Window size in bp

step_size <- 1	# Step size in bp

\# ----- Function to compute GC content -----

compute_gc_content <- function(seq, window_size, step_size) {

n <- length(seq)

num_windows <- floor((n - window_size) / step_size) + 1

gc_values <- numeric(num_windows)

for (i in seq(1, n - window_size + 1, by = step_size)) {

window <- subseq(seq, i, i + window_size - 1)

gc <- letterFrequency(window, letters = c("G", "C"), as.prob = TRUE)

gc_values[(i - 1) / step_size + 1] <- sum(gc)

}

return(gc_values)

}

\# ----- Perform GC content analysis -----

gc_content <- compute_gc_content(dna_seq, window_size, step_size)

avg_gc <- mean(gc_content)

\# ----- Prepare data frame for output and plot -----

gc_df <- data.frame(

Position = seq_along(gc_content),

GC_Content = gc_content

)

\# ----- Export to Excel -----

output_file <- paste0(marker_name, "_GCcontent.xlsx")

write_xlsx(gc_df, output_file)

cat("Results written to:", output_file, "\n")

\# ----- Define x-axis label -----

x_label <- paste0(marker_name, " (bp)")

\# ----- Visualization -----

plot <- ggplot(gc_df, aes(x = Position, y = GC_Content)) +

geom_point(size = 1) +

geom_hline(yintercept = avg_gc, linetype = "dashed", color = "black", size = 1) +

geom_smooth(method = "gam", formula = y ~ s(x, bs = "cs", k = 20)) +

scale_y_continuous(labels = scales::percent_format(accuracy = 1), limits = c(0, 1)) +

scale_x_continuous(breaks = seq(0, max(gc_df$Position), by = 100)) +

labs(

x = x_label,

y = "GC Content (%)"

) +

theme_minimal() +

theme(

axis.title = element_text(size = 14),

axis.text = element_text(size = 12),

plot.title = element_blank()

)

\# Display plot

print(plot)
