## Supplementary script 3 for "Incidence and attributes of chimeric COI and 18S sequences derived from nematode-infected arthropods"

\# Script: DNA stability analysis

\# Purpose: Perform a sliding window analysis of DNA stability using dinucleotide delta G values.

\# Load required libraries

library(Biostrings)

library(ggplot2)

library(writexl)

library(gam)

\# ----- Define genetic marker -----

marker_name <- "18S" # Change as needed (e.g., "COI", "ITS2")

\# ----- Define DNA sequence -----

\# Replace this string with the DNA sequence to analyze

dna_sequence <- DNAString("TCTAAGTACACACTGAACTAAAGTGAAACCGCGAAAGGCTCAGTATATCGGCTGTAATTCATTAGATCATATACTCAGTTACTTGGATAACTGTGGAAAATCTAGAGCTAATACATGCATTAATGCACGAACCGCAAGGAACGTGTGCAATTATTAGACAAACCAAGCGATGCTTGCATCGATTAACGGTGGAACTCTAGATAATTTTGCTGATCGTATGGTCTCGTACCGACGACAGATCTTTCAAATGTCTGCCCTATCAACTATTGATGGTAGTATAGAGGACTACCATGGTGGCAACGGGTAACGAGGAATCAGGGTTCGATTTCGGAGAGGCAGCATGAGAAACGGCTACCACATCTAAGGAAGGCAGCAGGCGCGTAAATTACCGATTTCCGGTTCGGAGAGGTAGTGACGAAAAATAACAATACAGGACTCATTAATGATGTCCTGTAATTGGAATGAGTTTAGTATAAATCCTTTAGCGAGGATCAAGTGGAGGGCAAGTCTGGTGCCAGCAGCCGCGGTAATTCCAGCTCCACTAGCGTATATTAAAATTGTTGCGGTTAAAACGTTCGAAGTTTGATTTTGTCCAGCACGGGTGTTACACCATAAGTATTGTGGTTGTAAATCACTCTTGATGCTAGGACTATAAGTTTGTGGCACCTGTGTAATGTCAAAGTTGCATAGGCAAAATGTGTATGGCGTGCCTTTAATCGGGTGTACCATATTGCATGCCCGACTATTTATTACCTTGAACAAATTAGAGTGCTCAAAGCAGGCTTTTATTGCCTTGAATAATCTTGCATGGAATAATGGAATAAGACCTCGATCAGTATATTTCATTGGTTTGAATTGATCATGAGGTAATGATTAATAAAAGCAGTTGGGGGCATTAGTATTACGACGCGAGAGGTGAAATTCATAGACCGTCGTAAGACTAACTGAAGAG")

\# ----- Define sliding window parameters -----

window_size <- 30

step_size <- 1

\# ----- Define dinucleotide delta G values -----

dinucleotide_scores <- c(

"GG" = -1.8, "CC" = -1.8, "GC" = -2.2, "CG" = -2.2,

"AA" = -1.0, "TT" = -1.0, "AT" = -0.9, "TA" = -0.6,

"GA" = -1.3, "AG" = -1.3, "GT" = -1.4, "TG" = -1.4,

"CA" = -1.4, "AC" = -1.4, "CT" = -1.3, "TC" = -1.3

)

\# ----- Define sliding window function -----

sliding_window_dinucleotide <- function(sequence, window_size, step_size) {

n <- length(sequence)

num_windows <- n - window_size + 1

delta_g <- numeric(num_windows)

for (i in 1:num_windows) {

window_seq <- as.character(substring(sequence, i, i + window_size - 1))

dinucs <- sapply(1:(window_size - 1), function(j) substr(window_seq, j, j + 1))

values <- sapply(dinucs, function(d) dinucleotide_scores[d])

delta_g[i] <- mean(values, na.rm = TRUE)

}

return(delta_g)

}

\# ----- Compute delta G values -----

dinucleotide_values <- sliding_window_dinucleotide(dna_sequence, window_size, step_size)

average_delta_g <- mean(dinucleotide_values, na.rm = TRUE)

\# ----- Prepare output data frame -----

data_dinucleotide <- data.frame(

Position = seq_along(dinucleotide_values),

DeltaG = dinucleotide_values

)

write_xlsx(data_dinucleotide, paste0(marker_name, "_dinucleotide_values.xlsx"))

\# ----- Define x-axis label -----

x_label <- paste0(marker_name, " (bp)")

\# ----- Plot results -----

ggplot(data_dinucleotide, aes(x = Position, y = DeltaG)) +

geom_point(size = 1) +

geom_hline(yintercept = average_delta_g, linetype = "dashed", color = "black", size = 1) +

geom_smooth(method = "gam", formula = y ~ s(x, bs = "cs", k = 20)) +

scale_y_continuous(limits = c(-2.5, 0), breaks = seq(-2.5, 0, 0.5)) +

scale_x_continuous(breaks = seq(0, length(dinucleotide_values), by = 100)) +

labs(

x = x_label,

y = expression(Delta * G^degree * " (kcal/mol)")

) +

theme_minimal() +

theme(

axis.title = element_text(size = 14),

axis.text = element_text(size = 12),

plot.title = element_blank()

)
