## Supplementary script 4 for "Incidence and attributes of chimeric COI and 18S sequences derived from nematode-infected arthropods"

\# Sliding window analysis of sequence identity

\# This script performs a sliding window analysis on sequence identity values.

\# The input file must be a CSV with two columns: 'Position' (numeric) indicating the alignment position, and 'Identity' (numeric) representing pairwise identity values at that position (ranging from 0 to 1).

\# Load required libraries

library(dplyr)

library(ggplot2)

\# Prompt user to select the identity CSV file

cat("Please select the identity CSV file...\n")

file_path <- file.choose()

\# Read the selected file

identity_data <- read.csv(file_path)

\# Define sliding window parameters

window_size <- 5

sliding_step <- 1

\# Initialize result container

num_rows <- nrow(identity_data)

results <- data.frame(position = numeric(0), average_identity = numeric(0))

\# Perform sliding window calculation

for (i in 1:(num_rows - window_size + 1)) {

start <- i

end <- i + window_size - 1

window_identity <- mean(identity_data$Identity[start:end])

results <- rbind(results, data.frame(position = identity_data$Position[start], average_identity = window_identity))

}

\# Plot results

ggplot(results, aes(x = position, y = average_identity)) +

geom_point() +

geom_smooth(method = "gam", formula = y ~ s(x, bs = "cs", k = 10)) +

labs(

x = "Alignment of 18S parent sequences (bp)",

y = "Identity (%)",

title = "Sliding window analysis of sequence identity"

) +

scale_x_continuous(breaks = seq(0, max(identity_data$Position), by = 100)) +

scale_y_continuous(limits = c(0, 1)) +

theme_minimal() +

theme(

axis.title = element_text(size = 14, color = "black"),

axis.text = element_text(size = 14, color = "black"),

plot.title = element_text(size = 16, hjust = 0.5)

)
